# The moon’s influence on the activity of tropical forest mammals

**DOI:** 10.1101/2024.02.20.581159

**Authors:** Richard Bischof, Andrea F. Vallejo-Vargas, Asunción Semper-Pascual, Simon D. Schowanek, Lydia Beaudrot, Daniel Turek, Patrick A. Jansen, Francesco Rovero, Steig E. Johnson, Marcela Guimarães Moreira Lima, Fernanda Santos, Eustrate Uzabaho, Santiago Espinosa, Jorge A. Ahumada, Robert Bitariho, Julia Salvador, Badru Mugerwa, Moses N. Sainge, Douglas Sheil

## Abstract

Changes in lunar illumination alter the balance of risks and opportunities for animals at night, influencing activity patterns and species interactions. Our knowledge about behavioral responses to moonlight is incomplete, yet it can serve to assess and predict how species respond to environmental changes such as light pollution or loss of canopy cover. As a baseline, we wish to examine if and how wildlife responds to the lunar cycle in some of the darkest places inhabited by terrestrial mammals: the floors of tropical forests.

We quantified the prevalence and direction of activity responses to the moon in tropical forest mammal communities. Using custom Bayesian multinomial logistic regression models, we analyzed long-term camera trapping data on 88 mammal species from 17 protected forests on three continents. We also tested the hypothesis that nocturnal species are more prone to avoiding moonlight, as well as quantified diel activity shifts in response to moonlight.

We found that, apparent avoidance of moonlight (lunar phobia, 16% of species) is more common than apparent attraction (lunar philia, 3% of species). The three species exhibiting lunar philia followed diurnal or diurnal-crepuscular activity patterns. Lunar phobia, detected in 14 species, is more pronounced with higher degree of nocturnality, and is disproportionately common among rodents. Strongly lunar phobic species were less active during moonlit nights, which in most cases also decreases their total daily activity.

Our findings indicate that moonlight influences animal behavior even beneath the forest canopy. This suggests that such impacts may be exacerbated in degraded and fragmented forests. Additionally, the effect of artificial light on wild communities is becoming increasingly apparent. Our study offers empirical data from protected tropical forests as a baseline for comparison with more disturbed areas, together with a robust approach for detecting activity shifts in response to environmental change.

**Open Research statement:** The data and code for performing the analyses described in this article are available at https://github.com/richbi/TropicalMoon.

## Introduction

The moon brightens the night. Changes in illumination associated with the 29-day lunar cycle alter the conditions faced by wildlife (Kronfeld-Schor et al. 2013). For some mammal species, especially those with limited night-vision or few nocturnal threats, the extra illumination provides periodic access to the night and associated foraging (Fernández-Duque et al. 2010, Prugh and Golden 2014) or travel opportunities (Gursky 2003). Other species are robbed of the cloak of darkness and become exposed to predators (Prugh and Golden 2014) or visible to prey (Pratas-Santiago et al. 2016).

The daily pattern of activity or “diel activity” of an individual, a population, or a species constitutes a fundamental part of its ecological niche and has been studied extensively (Bennie et al. 2014). Despite intuitive expectations for attraction to the moonlight (lunar philia) or avoidance of moonlight (lunar phobia) and accumulating evidence for each, our knowledge of wildlife responses to the moon and their prevalence in nature is still disjoint. While some species seem to respond strongly to lunar illumination (Kronfeld-Schor et al. 2013), others apparently do not respond at all (de Matos Dias et al. 2018, Zaman et al. 2022), for reasons not fully understood. The most comprehensive assessment of species responses to moonlight is a meta-analysis of 58 species that found that moonlight reduced the activity of species in open habitats yet increased the activity of species in forested environments (Prugh and Golden 2014).

However, this meta-analysis combined temperate forest species with tropical forest species. Moreover, the tropical forest species were predominately arboreal primates, which consistently responded positively to moonlight leaving open questions about other tropical forest mammals – including those living in the darkest part of forests - the forest floor. To date there has not yet been an assessment of the effects of moonlight on animal activity patterns based on community-level data collected across multiple locations and regions using standardized data collection and analytical methods.

There are both fundamental and applied reasons why we should identify responses to lunar phases and associated changes in illumination. First, the recurrent change in potential risks and opportunities faced by entire communities provides a testing ground for ecological theory about species adaptations (Bennie et al. 2014), interactions (Kronfeld-Schor et al. 2017), and the temporal dimension of the ecological niche (Kronfeld-Schor and Dayan 2003, Hut et al. 2012). Studies have tested for, and in some cases found, evidence that lunar illumination triggers niche shifting, with animals modifying when and where they are active dependent on the phase of the moon (Hut et al. 2012). Second, moonlight can serve as a model to help make predictions about the potential effect of artificial illumination – which is already impacting a substantial part of Earth (Cinzano et al. 2001, Falchi et al. 2016) – on wildlife behavior (Beier 2006, Rotics et al. 2011, Gilbert et al. 2023) and community dynamics (Meyer and Sullivan 2013, Gaston et al. 2014). Finally, knowledge about the relationship between illumination and animal behavior in densely canopied and less-impacted systems offers a baseline for detecting changes in human-modified habitats. Even natural light regimes change because of human-driven habitat alteration. For example, tropical forests, which harbor a substantial portion of earth’s biological diversity, are cleared, fragmented, and degraded at an alarming rate (Hansen et al. 2013, Pillay et al. 2022). Not only does this result in direct habitat modification, but also in reduced canopy cover which exposes forest-dwelling species to increased and prolonged solar and lunar illumination.

What is the prevalence and direction of responses to lunar phases in wildlife communities in some of the darkest places on earth, the floors of tropical forests? Do their inhabitants respond to moonlight like species in other environments and regions? Camera trapping offers an opportunity to answer these questions. If deployed long enough, camera traps record animal activity 24/7 throughout the lunar cycle and may thus capture responses of wildlife to changing levels of moonlight. We used images from a pantropical camera trap study in tropical forests across the globe. A standardized survey methodology allowed us to simultaneously examine diel and nocturnal activity of 88 mammal species spread over 16 orders and 35 families. Camera traps are now widely used for monitoring and studying terrestrial biodiversity (Burton et al. 2015, Steenweg et al. 2017, Semper-Pascual et al. 2022)and several studies have relied on time-stamped camera trap images to quantify and study animal diel activity (Rowcliffe et al. 2014, Frey et al. 2017, Vallejo-Vargas et al. 2022). We analyzed photographic detection data using a novel framework – multinomial regression combined with ternary classification – for consistent categorization and quantification of the temporal niche and shifts therein (see also Gerber et al. 2023). The flexible framework allowed us to not only compare levels of activity associated with different lunar phases, but also test hypotheses about how lunar illumination impacts activity beyond the night. Previous studies have shown that lunar illumination can trigger shifts in overall diel activity (Kronfeld-Schor et al. 2013). These changes may come about in different ways. At one extreme (fully additive), animals can reduce or increase their nocturnal activity during full moon, without a change in activity during day or twilight. This strategy will result in a corresponding decrease or increase in overall net activity. On the other extreme (fully compensatory), animals may shift activity into or out of the illuminated period, for example by moving their activity from the twilight period into the night, without a change in overall net activity.

The aim of this study was to better understand impacts of moonlight on animal activity. We investigated whether and how tropical forest mammals alter their diel activity in response to changing lunar phases. Specifically, we first assessed the prevalence of lunar philia and lunar phobia. Which species exhibit lunar philia or lunar phobia, and is one response to moonlight more prevalent than the other among mammals living under the dense canopy of tropical forests? Second, we tested for a link between a species’ degree of nocturnality and the response to lunar phases. Are nocturnal species more likely to manifest lunar phobia? Third, we quantified the extent to which mammals altered their diel activity in response to changes in lunar illumination. Do species responding to the different phases of the moon solely shift activity into or out of the night during moonlit periods without a change in overall activity levels (compensation) or do overall activity levels also change (additivity)?

## Methods

### Data collection

#### Camera trapping

We derived observations of mammal activity in protected tropical forests from camera trap data collected as part of the Tropical Ecology Assessment and Monitoring (TEAM) Network (Rovero and Ahumada 2017). Following a common protocol (Jansen et al. 2014), cameras were deployed between 2008 and 2017 throughout 17 protected areas in Indomalaya, the Neotropics, and the Afrotropics. The number of years of deployment varied between protected areas (2 years - 10 years; mean = 6.8 years), as did the number of locations sampled (60-90 camera trap locations; total: 1062). Spatial configuration and deployment were standardized, with cameras configured in either a 1x1km or 2x2km regular grid, at a height of approximately 30-50 centimeters off the ground. On average cameras were active for 33.2 days (SD=7.5). However, cameras were rotated sequentially until all sites were sampled within the wider sampling season. As a result, multiple lunar cycles are recorded at each protected area within a sampling season. For additional information about camera trapping protocols and species identification, see Rovero and Ahumada (2017). In this analysis, we included 2.1M photographic detections from 88 mammal species, i.e., species with ≥ 25 detections events during night (total across all protected areas; Supplementary Information Tables S1-S3). Due to apparent inconsistent identification from photographs, species in the genus *Tragulus* were considered jointly (*Tragulus sp.*)

### Analysis 1: Prevalence of lunar phobia and philia

#### Multinomial logistic regression

We use a Bayesian multinomial logistic regression model to simultaneously assess diel (entire 24-hour period) and nocturnal (lunar) activity patterns. We distinguished three diel periods (day, night, and twilight) and three lunar periods (full moon, transitional, new moon). We chose discrete diel and lunar periods (see definitions below) instead of continuous values based on illumination (Smielak 2023), as it enabled the multinomial analysis and an intuitive categorization of activity (Figs 1 and 2).

**Figure 1.**
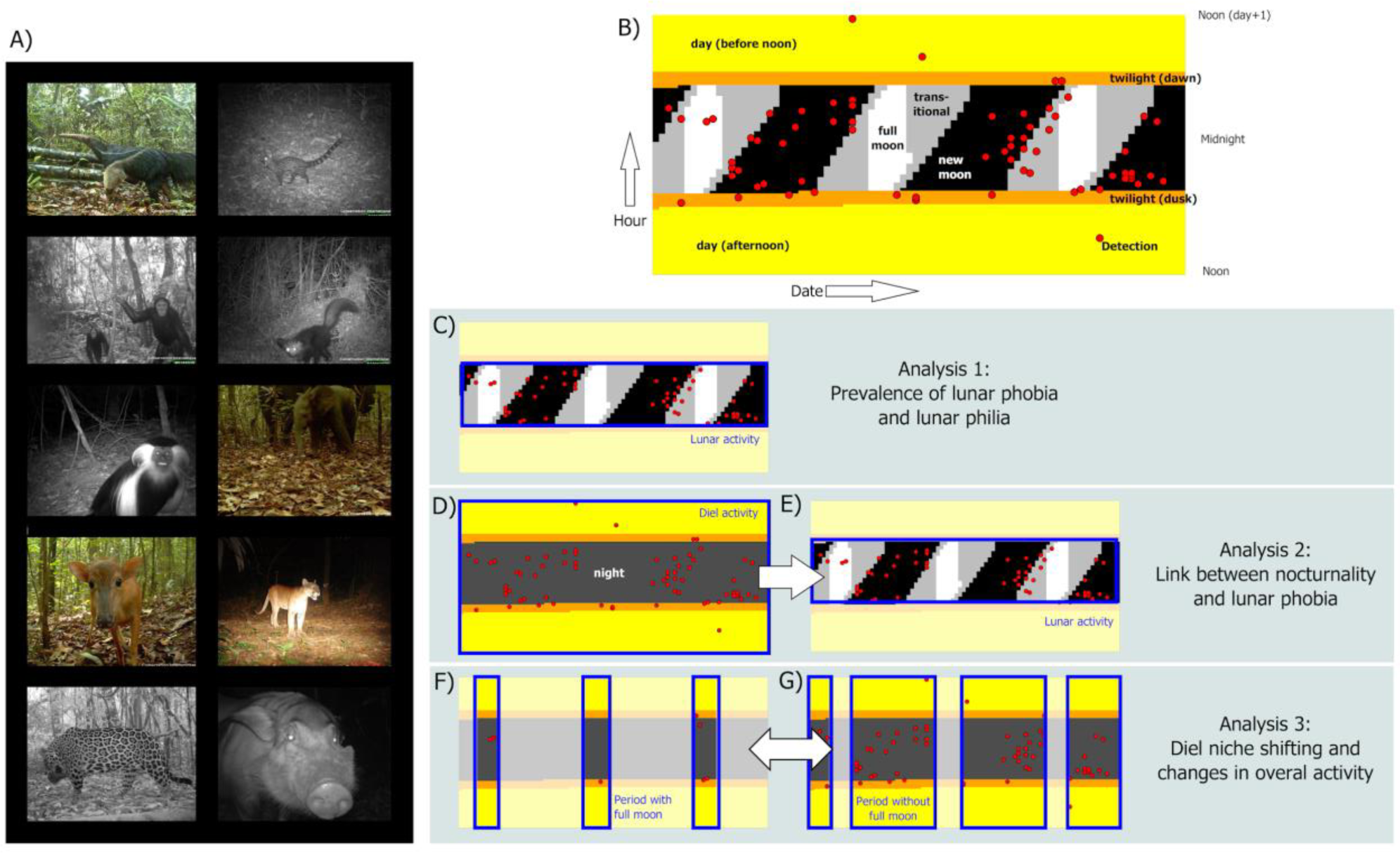
Illustration of the study design. Time-stamped camera trap images (A) are aggregated into 15- minute intervals and mapped onto available site-specific diel and lunar periods (B). Red dots in the example belong to the nine-banded armadillo (*Dasypus novemcinctus*), an apparently lunar phobic species. Multinomial logistic regression models are used to quantify the probability of a species using a given diel or lunar period. Three analyses explore 1) the prevalence of lunar phobia and lunar philia during nocturnal activity (C), 2) the effect that the level of nocturnality of a species (D) has on its propensity to exhibit lunar phobia (E), and 3) changes in diel activity patterns and total activity levels during periods with full moon (F) vs. other lunar phases (G). The blue boxes delineate the part of the diel region involved in each assessment or comparison. Photos: TEAM network.

**Figure 2.**
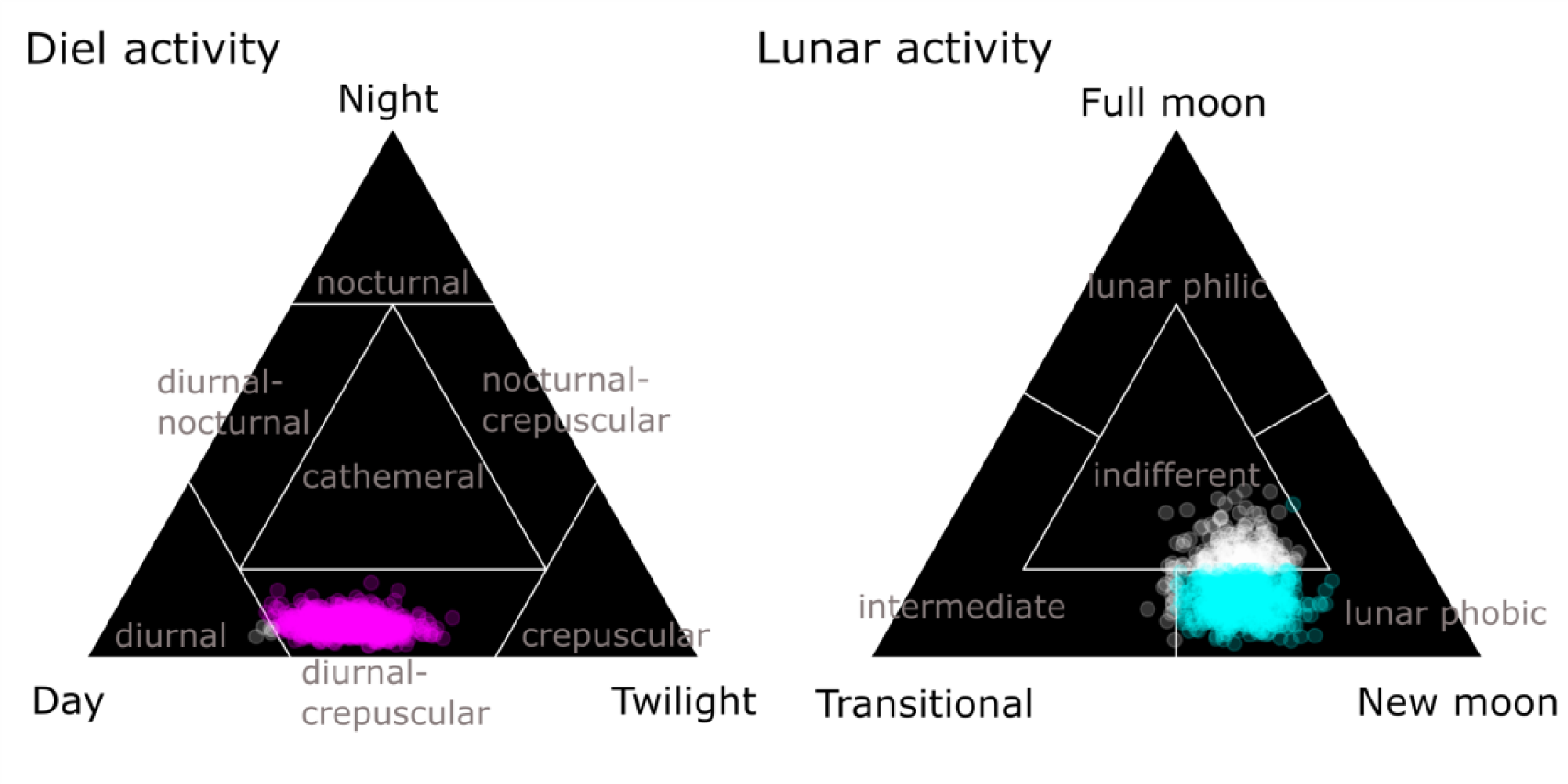
Posterior samples (dots) of multinomial probabilities mapped onto ternary diagrams. Shown are example posterior samples for diel activity (left; activity during day, night, and twilight periods) and lunar activity (right; nocturnal activity during full moon, new moon, and transitional phases). The ternary diagrams are divided into seven and four regions for diel and lunar activity delineation respectively. Designation to activity categories (grey text) is made according to the region into which the majority (colored dots) of posterior samples are mapped. The examples show a diurnal-crepuscular (left) and a lunar phobic species (right).

This model contained two submodels, one for diel activity and one for lunar activity. The submodel for diel activity consisted of a multinomial logistic regression model to estimate species-specific probability of photographic capture in one of the three major diel periods (day, night, twilight; see also Gallo et al. 2022):

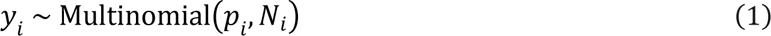

Here, *Y*_*i*_ is the length-3 vector of the number of independent photographic capture events of species *i* in each diel period, *N*_*i*_ the total number of detections (*N*_*i*_ = ∑ *Y*_*i*_) of that species, and *p*_*i*_ the length-3 vector of probabilities of detection in each diel period.

The multinomial probability vector can be defined using logistic regression:

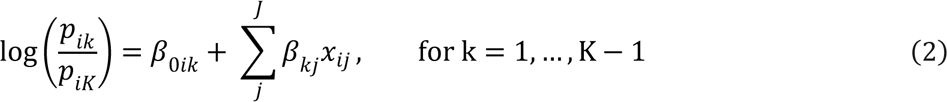

where *β*_*i*0*k*_ is the species-specific intercept term associated with categorical outcome *k* (diel period) out of the total possible number of outcomes *K* (i.e., 3: day, night, twilight), *β*_*kj*_ the *j*’th out of a set of *J* coefficients associated with predictor *x*_*ij*_. The quotient on the left side of eq 2 signifies that the last outcome (*p*_*iK*_) serves as a reference value for the other *K* − 1 outcomes (*p*_*ik*_ ).

Predictor variables and associated coefficients shown in eq 2, were omitted in our multinomial logistic regression model for diel activity as we were primarily interested in estimating intercepts and corresponding probabilities:

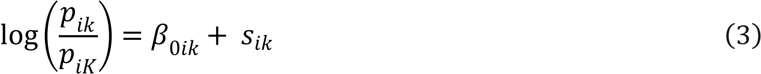

In addition to species-specific intercepts, we incorporated an offset variable *s*_*ik*_ defined as the proportion of time (rounded to number of hours in our analysis) during which cameras were active (available for making photographic captures) within each diel period *k*, relative to the reference period *K*. The offset variable serves the purpose to account for differences in “availability” (see also Gallo et al. 2022), and has the effect of adjusting the estimated intercept according to the amount of camera trap effort in each observational record. For example, the crepuscular period is significantly shorter than periods of daylight and night. Similarly, the period of full moon only makes up a small proportion of total nighttime (Figure 1). The relative “availability” of different diel periods also changes over the course of seasons, particularly at latitudes farther from the equator. The model thus produces comparable estimates of selection for or against a given period, reflecting “density” of activity (hours with photographic captures per total hours of camera operation during a period) rather than pure activity volume.

The probabilities of interest are thus:

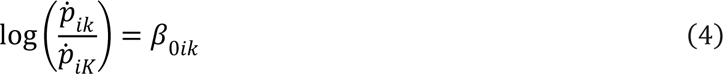

Where both *ṗ*_*ik*_ and *p*_*ik*_ are scaled to sum to 1 across the *K* multinomial outcomes.

Detection data (*Y*_*i*_) used in the analysis constituted the number of hours with at least one detection of a given species in each diel period (daylight, twilight, and night) at each camera trap site, summed across all sites. The sum of the period-specific activity of a species makes up its total activity *N*_*i*_. Availability (to calculate the offset *s*_*k*_) was derived as the number of hours that fell into a given diel period at each camera trap site, summed across all camera trap sites. Diel periods were delineated using local (study area specific) astronomical sunrise, sunset and twilight times, assuming a flat landscape and obtained using R package ‘suncalc’ (Thieurmel and Elmarhraoui 2022). Dawn was delineated by the beginning of astronomical twilight (sun 18° below the theoretical horizon) and sunrise (when the bottom edge of the sun touches the theoretical horizon). Dusk was delineated by the beginning of sunset and astronomical sunset (sun 18° below the horizon). Night was delineated as the period between astronomical dusk and dawn and day as the period between sunrise and sunset.

The submodel for lunar activity was structurally identical to the diel activity model described above. In the lunar submodel, the three multinomial probabilities (*ṗ*_*ik*_ ) represent species-specific estimates of the probability of photographic capture in one of the three major lunar periods, roughly corresponding to full moon, new moon, and the combined intermediate phases. Moon phases were delineated for the night (as defined above) using moon altitude (angular elevation) and illumination, again with R package ‘suncalc’. Full moon was defined as the period when the moon had an altitude ≥ 18° above the theoretical horizon and was ≥90% illuminated. New moon was defined as the period when the moon had an altitude < 18° above the theoretical horizon or was <10% illuminated. All other nocturnal periods were designated as transitional phases.

#### Model fitting

We fitted multinomial models using Markov chain Monte Carlo (MCMC) simulation with NIMBLE version 1.0.0 (de Valpine et al. 2017) in R version 4.3.0 (Team 2023). We ran 4 chains with 40000 iterations each, including a 20000-iterations burn-in period. Chains were thinned by a factor of 5. We considered models as converged when the Gelman-Rubin diagnostics (Gelman 1996)was ≤ 1.1 for all parameters and after visually inspecting trace plots.

#### Designation of diel and lunar activity categories

For visual inspection, categorization, and presentation, species-specific posterior samples of multinomial probabilities produced by the Markov chain Monte Carlo MCMC analysis were plotted onto Ternary diagrams (Shepard 1954) using package ‘ternary’ (Smith 2017) in R. The diel and lunar activity pattern of each species was delineated with the help of the ternary diagrams. We considered several alternative ternary configurations for categorization (Shepard 1954, Schlee 1973, Santini et al. 2005, Nakamura et al. 2018), but ultimately opted for a subdivision into 7 regions for diel activity and 4 regions for nocturnal activity as it relates to the lunar cycle. The lower number of categories for nocturnal activity was motivated by the lower sample size (only observations made during the night are considered for categorizing lunar activity) and ease of interpretation. For categorizing diel activity, we divided the ternary diagram into these 7 regions (Fig. 2): 3 corner triangles (each capturing cases that contain >2/6 of all activity) for the “pure” diel activity categories (e.g., diurnal), three transitional regions between pairs of corner regions for intermediate categories (e.g., diurnal-crepuscular), and one central triangle that indicates cathemerality (activity during all diel periods). This classification follows Shepard’s (1954) approach for delineating soil categories, but without the additional splitting of the intermediate regions along the sides of the ternary. We divided the ternary diagram for lunar activity categorization into only these 4 regions (Fig. 2): one central triangle (identical to the cathemeral region in the ternary diagram for diel activity) representing indifference (referred to as “neutrality” by Gursky 2003) to the phase of the moon and 3 main lunar categories (activity during full moon, new moon, and intermediate lunar phases).

Species activity is categorized based on the position of the posterior distribution of multinomial probabilities (*ṗ*_*k*_) within the ternary space. As a species-level designation (diel and lunar activity category/strategy/niche), we based designation on the region that contained the majority of the posterior samples of the multinomial hyperparameter, depending on which one of the seven (diel) and four (lunar) regions the majority of posterior samples fell into.

### Analysis 2: Link between nocturnality and lunar phobia

To estimate the relationship between diel and lunar activity we used the model from Analysis 1 as a starting point, but now linking the lunar activity submodel with the diel activity submodel:

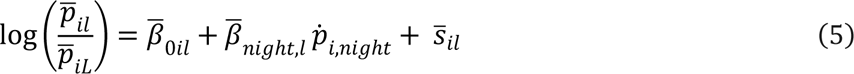

Where *ṗ*_*i,night*_ is the strength of selection for nocturnal activity estimated in the dial submodel and *β*_*night,i*_ is its effect on the multinomial probability associated with lunar period *l* out of a total of *L* lunar periods. Whereas *ṗ*_*i,night*_ is species specific, we estimate one coefficient *β̄*_*night,i*_ in this analysis, across the entire species data set. Model fitting proceeded as in Analysis 1. We used the posterior distribution of *β̇*_*night, i*_ and species-specific *β̄*_*oil*_ to derive fitted values (means) and associated 95% Bayesian credible intervals (BCI) of the link between nocturnality and the probability of association with new moon and full moon periods.

### Analysis 3: Diel activity shifting and changes in overall activity levels

We used a third Bayesian model to assess whether and how animals altered their diel activity in response to changes in lunar illumination (Fig. 1). Specifically, we tested whether species categorized as lunar phobic during the first analysis 1) reduced their overall activity (number of photographic capture events) during the periods (Fig. 1F-G) that contained nights with full moon and/or 2) shifted their diel activity towards daylight or twilight. Conversely, for species categorized as lunar philic, we tested whether they 1) increased their overall activity during the multi-day time periods containing bright nights and/or 2) shifted their diel activity towards the night.

We used two submodels, one for modelling the number of photographic detection events during 24-hour periods with and without at least one hour of full moon at night and a multinomial logistic model for overall diel activity during the same time periods.

The model for the total number of photographic detection events *n*_*i*_ for species *i* during a given period (days with vs without full moon at night) was formulated as a generalized linear model with a log-link (Poisson regression):

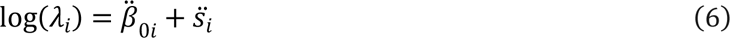

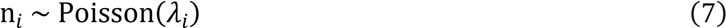

Where λ_*i*_ is the parameter of the Poisson distribution (expected number of events) and *β̈*_0*i*_ the species- specific intercept. As in the multinomial models (equations 3 and 5), we included an offset term *s̈*^*i*^ to account for differences in availability, provided as the total operational camera trap hours associated with a given lunar phase over all camera trap sites and sampling seasons in protected areas where species *i* was detected at least once. This allowed direct comparison of periods with and without moonlit nights via *λ̇*_*i*_, derived as

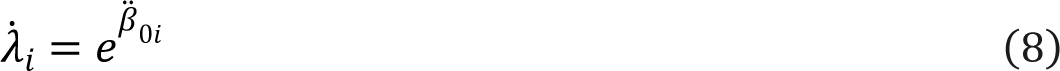

The multinomial model for diel activity was identical to the model defined with equations 1 and 3. The main difference between the diel activity model in analyses 1, 2, and 3 was in the design: whereas in analyses 1 and 2 we estimated the multinomial probabilities of being active during the three diel periods (day, night, twilight) at any point during monitoring, in analysis 3 we estimated separate multinomial probabilities for periods (multiple days) with and without at least 1 hour of full moon at night (Fig. 1).

Model fitting and assessment of convergence/mixing was performed as in analysis 1 and 2. Diel activity was categorized using the ternary approach described earlier. Changes in diel activity during periods with full moon that resulted in a change of activity category were considered evidence of temporal niche switching. Changes in diel activity without a change in activity category but a difference in the posteriors for *ṗ*_*night*_ associated with new moon vs. all other phases whose 95% BCI did not include zero were considered activity timing shift, a term also used in (Gilbert et al. 2023). We subtracted the posterior sample for λ̇_*i*_ for periods without moonlit nights from those with to derive species-specific posteriors of the effect of full moon on overall activity levels. We considered a species to show evidence of altered overall activity levels in response to lunar illumination when the 95% BCI of this derived variable did not include zero.

## Results

### Diel activity and overview

Of the 88 species included in the analysis, we categorized 20 species as predominantly nocturnal and nine as diurnal, following the multinomial regression analysis controlling for temporal availability and the ternary classification scheme (Fig. 2, Supplementary Information Tables S1-S3). Only one species (common tapeti, *Sylvilagus brasiliensis*) was categorized as predominantly crepuscular. Most species (42) fell into one of the two intermediate categories involving crepuscularity (Fig. 3). All remaining species (16) were categorized as cathemeral (Supplementary Information Tables S1-S3). Cathemeral designation, by nature of its position within the ternary, is associated with greater uncertainty (Gerber et al. 2024). In data-sparse situations it may be difficult to distinguish between a species being truly cathemeral and the model not having enough information to assign the species to another category. However, all species categorized as cathemeral in this analysis had more than 100 observations (hours with at least one detection; mean = 770, range = 136– 3250; Supplementary Information Tables S1-S3).

**Figure 3.**
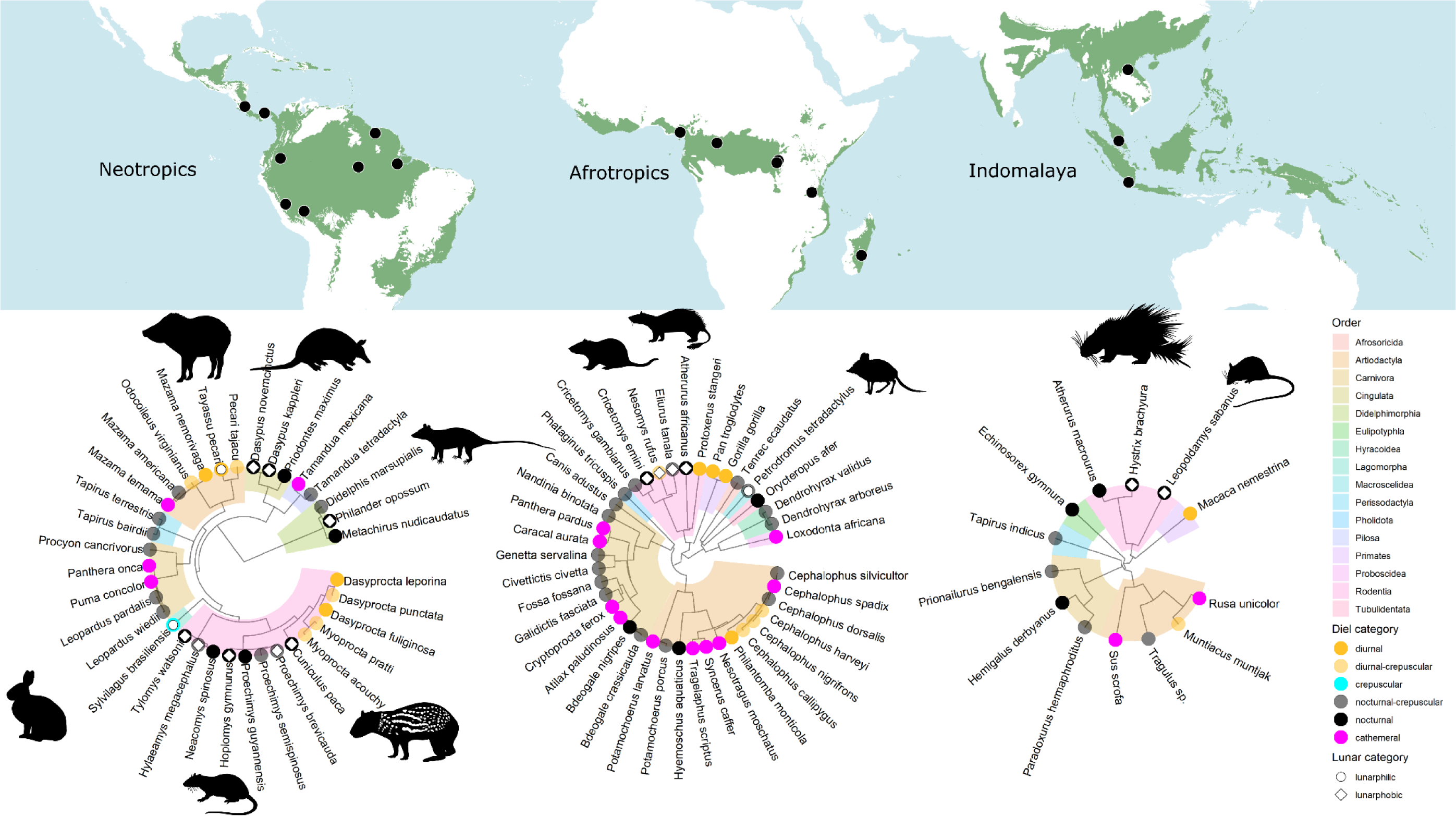
Diel and lunar categorization charted across phylogenetic trees of tropical forest mammals in three realms. The distribution of tropical moist broadleaf forest (green regions) and location of study areas (black dots) included in this analysis are shown on the map. Lunar category is only indicated for species that were unambiguously designated as either lunar phobic or lunar philic. Lunar phobia, manifested as reduced activity during moonlit nights, was more common than lunar philia, increased activity during moonlit nights. Rodents, particularly nocturnal species, were overrepresented among lunar phobic species, followed by members of the Cingulata (including armadillos) and Didelphimorphia (opossums). Phylopic silhouettes credits: *Cuniculus paca, Dasypus novemcinctus, Silvilagus brasiliensis* and *Tayassu pecari* by Gabriela Palomo-Munoz*; Philander opossum* by Milena Cavalcanti, Patricia Pilatti & Diego Astúa; *Hoplomys gymurus, Atherurus africanus, Cricetomys emini, Hystrix brachyura* and *Leopoldamys sabanus* provided by Andrea F. Vallejo Vargas; *Petrodromus tetradactylus* under universal public domain license.

**Figure 4.**
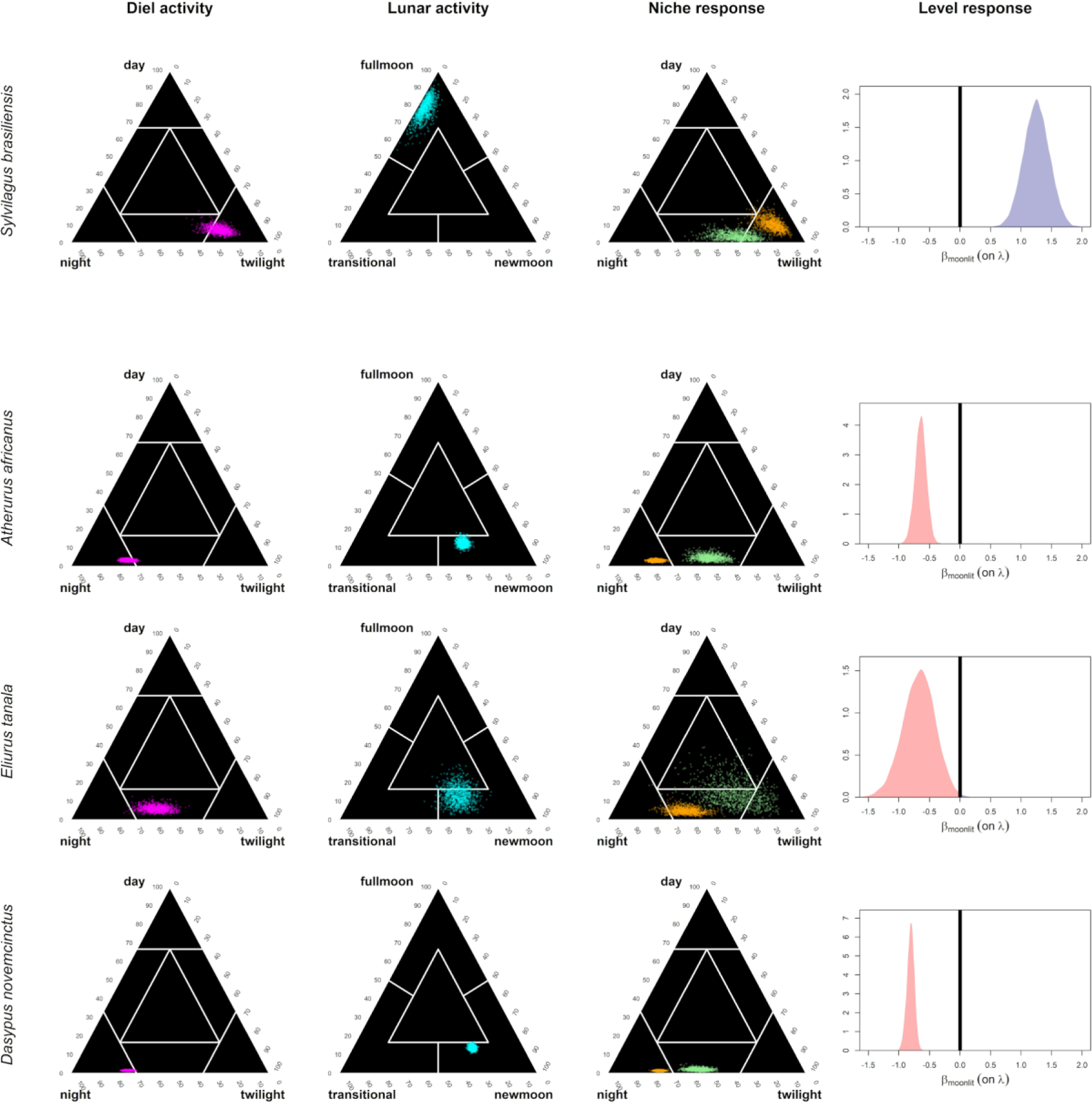
Overview of ternary classifications for diel and lunar activity (columns 1-2) and responses to lunar phases (column 3 and 4) for four example species (rows). The species shown in the top row (common tapeti or forest cottontail) was classified as lunar philic, whereas the bottom three rows show lunar phobic species. Column 1: ternary plots of diel activity posteriors. Column 2: ternary plot of lunar activity posteriors. Column 3: ternary plots showing difference of diel activity (potential temporal niche shifting) between periods with full moon (green) and without moonlit nights (orange). Column 4: posterior distribution of the difference between overall activity (related to the number of photographic detection events) during periods with vs. without full moon. Negative (red) and positive (blue) values indicate a reduction or an increase, respectively, in overall activity during periods (multiple 24-hour periods, Fig. 1) with full moon. See Supplementary Information Figures S1-S3 for results for all species classified as either lunar phobic or philic.

### Prevalence of lunar phobia and philia

Of the 88 species included in the analysis, 14 were categorized as lunar phobic and three as lunar philic (Fig. 3, Supplementary Information Tables S1-S3). Only one species (Forest giant squirrel, *Protoxerus stangeri*) was categorized as selecting for intermediate lunar phases (“transitional”). Rodents were the most common lunar phobic species (11), followed by armadillos (2), and one opossum (gray four-eyed opossum, *Philander opossum*). The representation of rodents among lunar phobic species (79%) was disproportional to their prevalence (25%) among the species in our sample. The three mammal species exhibiting lunar philia were the white-lipped peccary (*Tayassu pecari*, order Artiodactyla) and the common tapeti (order Lagomorpha) in the Neotropics, and the four-toed elephant shrew (*Petrodromus tetradactylus*, order Macroscelidea) in the Afrotropics. The remaining 70 species were categorized as indifferent towards lunar phases, either because their nocturnal activity was not impacted by lunar illumination or because their data had such a high noise-to signal ratio that it prevented designation to one of the peripheral ternary regions (Supplementary Information Tables S1-S3). In our dataset, 14 (20%) of the species categorized as indifferent towards lunar phases had less than 50 observations during the night and we consider these species as data-sparse. Nonetheless the sample size was relatively high with an average of 445 nocturnal observations (range: 25 – 4189; Supplementary Information Tables S1-S3) of species categorized as indifferent towards lunar phases.

### Link between nocturnality and lunar phobia

Species with a greater probability of being active at night were more likely to be more active also at new moon (*β*_*night.new moon*_ = 1.55, 95% CrI: 1.46 to 1.64, Fig. 5) and, conversely, less likely to be active at full moon (*β*_*night.full moon*_ = -1.12, 95% CrI: -1.25 to -0.99, Fig. 5). This effect was also reflected in our categorization of diel activity of species identified as lunar phobic. Thirteen of the fourteen species categorized as lunar phobic also exhibited a nocturnal or nocturnal-crepuscular diel activity pattern, and only one was categorized as diurnal-crepuscular (Fig. 3). Of the lunar philic species, one was diurnal, one diurnal-crepuscular, and one nocturnal-crepuscular (Fig. 3).

**Figure 5.**
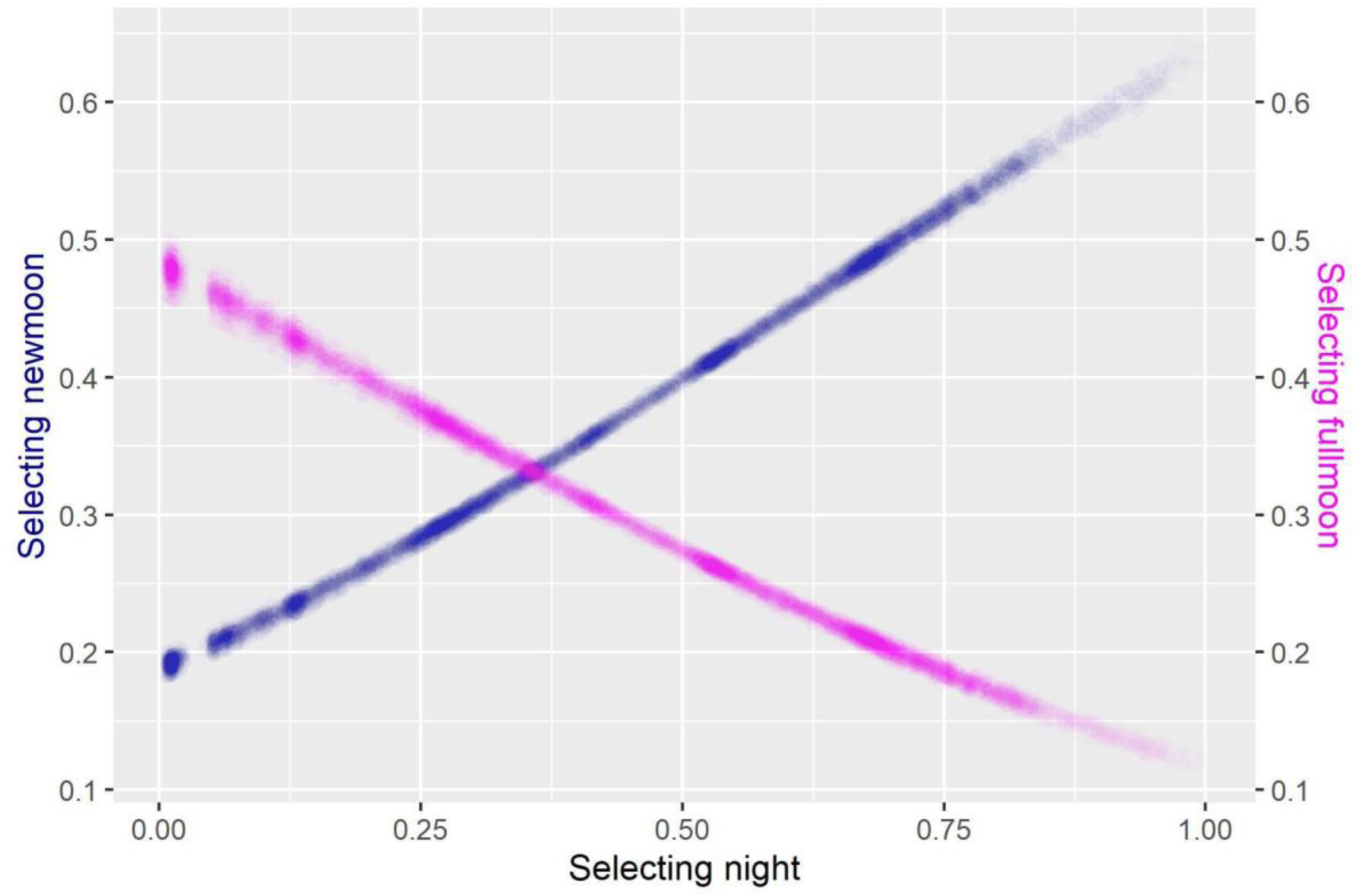
Effect of the probability of nocturnal activity on the probability of activity during new moon (blue) and full moon (magenta). The plot shows posterior predictions, with the intensity of shading (opaqueness) corresponding to the posterior distribution of *ṗ*_*night*_(x-axis, eq 3), *ṗ*_*full moon*_ (left y-axis) , and *ṗ*_*new moon*_ (right y-axis) across all 88 species included in the analysis.

### Temporal niche shifting and changes in overall activity

Eleven of the 14 lunar phobic species significantly reduced their overall activity level during periods with moonlit nights and eight of these shifted their diel behavior to become less nocturnal (Figs 3 and 5; Supplementary Information Fig. S2). Three species classified as lunar phobic (*Philander opossum, Hoplomys gymnurus, Tylomys watsoni*) reduced their overall activity without a significant shift in diel behavior, and two species (*Nesomys rufus, Protoxerus stangeri*) shifted their diel behavior to be less nocturnal without a significant reduction in overall activity (Fig. 6). Of the three species identified as lunar philic, one (*Sylvilagus basiliensis*) shifted to more nocturnal activity and increased its overall activity, one (*Tayassu pecari*) only shifted its activity to become more nocturnal, and one (*Petrodomus tetradactylus*) only increased its overall activity during periods with moonlit nights (Fig. 6). Following our categorization of diel behavior, 9 of the 14 lunar phobic species switched their temporal niche from nocturnal to nocturnal-crepuscular during periods with full moon. See Supplementary Information Figures S1 and S2 for detailed results for all species classified as lunar phobic or philic.

**Figure 6.**
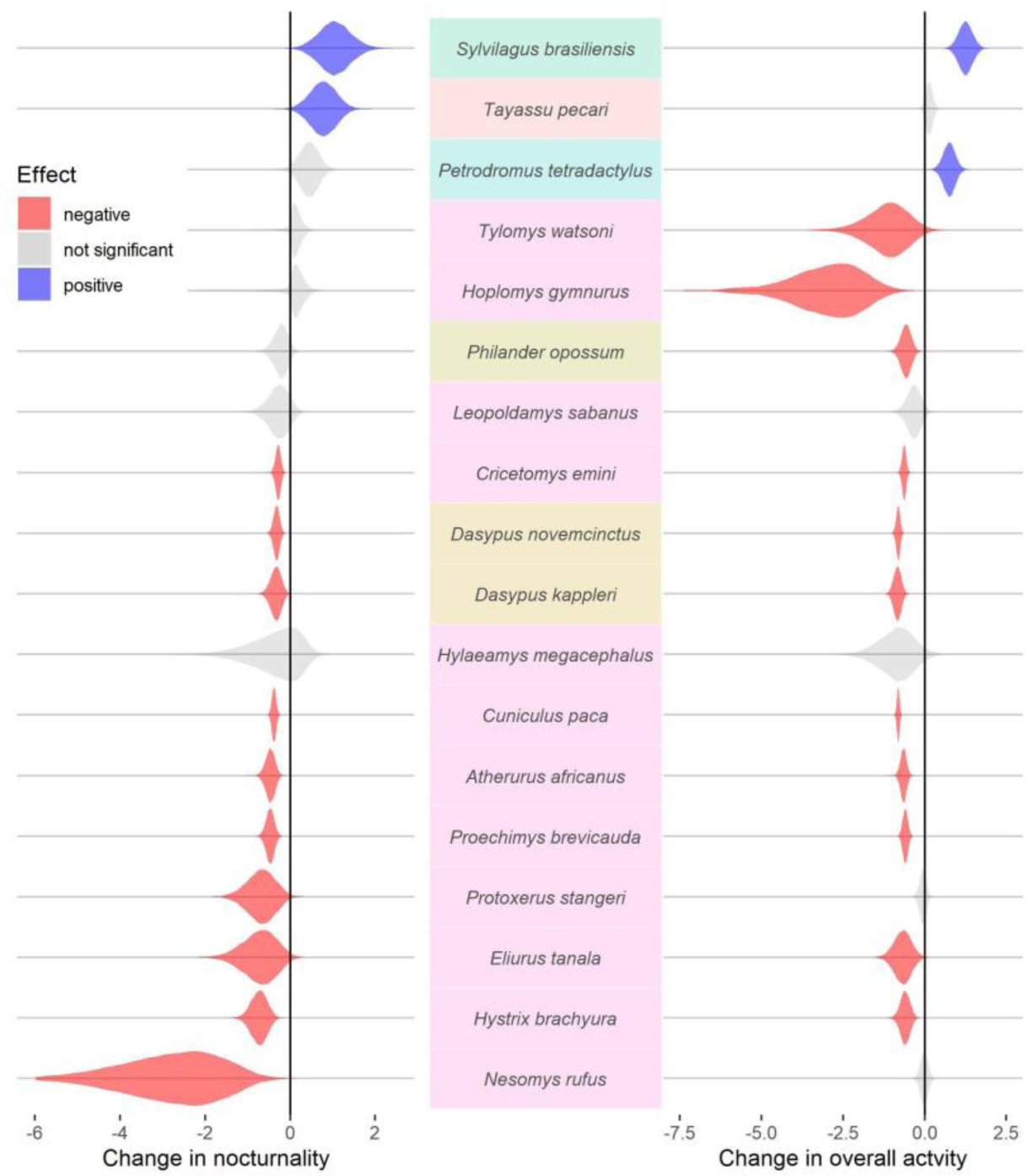
Changes in nocturnality (left) and overall diel activity levels (right) of forest mammals during periods with and without full moon. Responses (“Effect”) are represented by the posterior distributions of coefficient estimates shown as violins. Species names are shown on color-coded backgrounds according to the order they belong to (see Fig. 3 for key). The top 3 species were categorized as lunar philic, the remainder as lunar phobic.

## Discussion

Wildlife responses to moon phases are still poorly understood. We applied a novel analysis of activity patterns to data from standardized camera trapping in 17 tropical forests across the globe. We found that even in the understory of protected tropical forests, characterized by densely shaded habitats, the moon’s phases impact the activity of mammal species. Lunar phobia was more common than lunar philia, with rodents being the most common lunar phobic taxonomic group. Additionally, we found that nocturnal species are more active during new moon than during other lunar phases. Finally, our quantitative study revealed that species may avoid bright moonlight by shifting their activity towards other parts of the diel cycle, by reducing overall activity, or both. These findings indicate that changes in illumination (e.g., through deforestation or artificial illumination) could affect species activity, and ultimately interactions in tropical forest communities.

### Lunar philia and lunar phobia in tropical forest mammal communities

Unlike prior meta-analysis results showing that forest dwelling species increased activity during moonlight (Prugh and Golden 2014), our standardized assessment shows that the most common tropical terrestrial mammal response to moonlight is reduced activity. Lunar philia is rare among terrestrial tropical forest mammals. Only three species, among the 88 species studied here, significantly increased their exposure to camera traps during hours filled with moonlight (Fig. 3). Lunar philia has previously been reported as comparatively rare and has been associated with species, such as arboreal primates, relying on visual cues for foraging and predator avoidance (Prugh and Golden 2014). The only species classified as lunar philic in our study were a peccary (white-lipped peccary, *Tayassu pecari*), a rabbit (common tapeti, *Silvilagus brasiliensis*), and an elephant shrew (four-toed elephant shrew, *Petrodromus tetradactylus*). Apparent lunar philia in the common tapeti has previously been reported in Argentina (Huck et al. 2017), and contrasts lunar phobic behavior reported in another lagomorph, the snowshoe hare (*Lepus americanus*, Griffin et al. 2005). Lunar philia in elephant shrews is consistent with descriptive studies on the order (Woodall et al. 1989). Similarly, white-lipped peccary have been reported to change routes and increase movement in the forest during full moon (Serrano et al. 2010, Hernández 2015). *T. pecari* is a large group-living mammal (40 kg) which may make it less vulnerable predators.

Lunar phobia, in contrast, was common. It was exhibited primarily by small-to-medium sized mammals that are prey of carnivores such as ocelots (*Leopardus pardalis*) or jaguars (*Panthera onca*, Moreno et al. 2006, Pratas-Santiago et al. 2016). Nevertheless, the fact that lunar phobia was disproportionately found in rodents, suggests that there are evolutionary differences among prey groups that influence responses to moonlight above and beyond ecological responses to predators as prey, potentially related to their sensory ecology. The paca, one of the largest lunar phobic rodents in our study (8000g) has been classified as lunar phobic in other study areas (but see Michalski and Norris 2011), as were armadillos (Harmsen et al. 2011, Pratas-Santiago et al. 2017). Pacas and armadillos need to avoid natural predators but also hunting by rural and indigenous communities (Redford and Robinson 1987, Pires Mesquita et al. 2018). Although Harmsen et al. (2011) detected no changes in the activity of the common opossum (*Didelphis marsupialis*) in response to moonlight, evidence on lunar phobia in the gray four-eyed opossum (*Philander opossum*) in our study matches findings for members of the Didelphimorphia order reported in other studies (e.g., *Didelphis aurita*, *Calouromys philander*; Julien-Laferrière 1997, Tripodi et al. 2023).

We simultaneously analyzed activity data from many species using one standardized approach. Across that diverse community of mammals, some species were observed more frequently than others.

Technically, all activity data, no matter how sparse, can be designated to one of the categories represented by the areas delineated in the ternary diagram. However, observation of activity is imperfect in nature, either because not all individuals are observed or individuals are not observed all of the time, or both. The probability of making an erroneous designation (i.e., an activity category other than the true one) is liable to increase as sample size decreases. Designation to the cathemeral (diel activity) and indifferent (lunar activity) categories, by nature of their position within the ternary, are probably the most error prone and most vulnerable to data paucity. Determination of cathemerality was less ambiguous, as all species in this study had more than 100 observations for diel activity categorization, with an average of 796 observations per species. However, 22% and 44% of species categorized indifferent to lunar phases had < 50 and < 100 hourly observations respectively. At least for these, we cannot reliably distinguish true indifference to lunar phases from an insufficient sample size for making a designation.

### The link between lunar phobia and nocturnality

The more nocturnal species are, the more likely they are to exhibit lunar phobic behavior (Fig. 5). Lunar phobia has been explained as a behavioral adaptation by nocturnal species, intended to avoid the elevated predation risk during periods with higher illumination (Daly et al. 1992, Kronfeld-Schor et al. 2013). The large number of species included in our study and a standardized classification approach yielded support for this explanation. The avoidance of moonlight could reduce vulnerability to detection by visually-hunting predators or, in the case of lunar phobic predators, detection by prey. Rodents, for example, generally seem to reduce foraging activity during bright nights (Price et al. 1984, Longland and Price 1991, Prugh and Golden 2014). Conversely, we found that lower nocturnality was associated with higher activity during full moon periods (Fig. 5). The three species identified as lunar philic were diurnal, crepuscular, and crepuscular-nocturnal. Two of these species significantly increased their overall level of activity during moonlit periods (Fig. 6). Meanwhile, 13 of the 14 lunar phobic species were nocturnal or nocturnal-crepuscular, and only one was categorized as diurnal-crepuscular (Fig. 3). Thus, moonlight appears to give species adapted to daylight and twilight better visual access to the night.

### Temporal niche shifting in response to moon phases

Lunar illumination can trigger changes in diel activity. We found that during periods with full moon, eight species (all classified as lunar phobic) changed their diel activity from nocturnal to crepuscular-nocturnal. These species also decreased their overall activity level. These results are in line with observations on snowshoe hares (*Lepus americanus*, Studd et al. 2019) and Marriam’s kangaroo rats (*Dipodomys merriani*, Daly et al. 1992), which reduce their activity during moonlit nights and increase their diurnal and crepuscular activity, respectively. Potentially indicative of compensatory response to lunar illumination, two species reduced their nocturnal activity without an apparent reduction on overall activity during periods with full moon. Conversely, three species reduced their overall activity without a clear shift in nocturnality, suggesting a response closer to the additive end of the spectrum. One of the three lunar philic species increased its overall activity levels during periods with illuminated nights and shifted from diurnal or crepuscular towards more nocturnal activity. This echoes the findings from studies on owl monkeys (*Aotus azarai*), a generally nocturnal species, which increases its activity during full moon, and, during new moon, shifts its activity from night towards day (Erkert 2008, Fernández-Duque et al. 2010). Regardless of the strategy chosen in response to changing lunar illumination, our results showed evidence of temporal niche shifting. Two lunar phobic species in our study exhibited a reduction in overall activity levels without a noticeable shift towards crepuscular behavior. We could speculate that this is a result of behavioral inflexibility (i.e., strict nocturnality); although it may be purely a result of insufficient statistical power or reflect a shift to habitat strata less well-covered by camera traps.

### Methodological insights and other considerations

In this study, we adjusted and deployed a novel framework to delineate diel and nocturnal activity categories using multinomial probability distributions, and ternary diagrams (Gallo et al. 2022, Gerber et al. 2024). This approach is both visually intuitive and quantitative, facilitating detection of ecological patterns related to activity, such as temporal niche partitioning and niche shifting/switching in response to moonlight. Any analytical approach that can estimate the probability of designation and the associated uncertainty can be substituted for the Bayesian multinomial approach used here. The advantage of the latter is that it produces posterior samples of multinomial probabilities, which readily allow propagation of uncertainty to the ternary projection and subsequent classification.

We recorded species’ diel activity detectable with camera traps. Camera trap data lend themselves to comparative and comprehensive diel activity studies as they monitor entire communities (Cid et al. 2020, Vallejo-Vargas et al. 2022) and are non-invasive, or at least less invasive than traditional methods such as direct observation and telemetry. The rapidly expanding spatial and temporal scope of camera trapping in wildlife ecology offers opportunities for revising and filling gaps in our understanding of the temporal niche of wildlife and its dynamics. Nonetheless, camera trapping has limitations and inferences should be drawn with caution. For example, if arboreal or scansorial animals shift their activity to lower forest strata during moonlit periods or if species move into more densely vegetated areas from beyond forest edges, lunar phobia may increase terrestrial activity as detected through photographic captures by understory cameras. In our study, however, all but one of the species with lunar responses are classified as terrestrial (Wilman et al. 2014). Yet, other sampling methods (or sampling in other strata; Bowler et al. 2017, Haysom et al. 2021) may in some situations and for certain species be more suitable to obtain reliable data on activity. Any sampling approach that does not influence activity itself and produces timestamped observations can be used.

### Implications

The influence of natural and artificial light is an increasingly important topic in wildlife conservation and ecosystem functioning (Gaston et al. 2017, Hirt et al. 2023). Yet we still know too little about the implications of artificial light on the activity of mammals (Hoffmann et al. 2022). The prevalence of lunar phobia in our study suggests there may be more losers than winners when illumination increases in tropical forests. Moreover, most lunar phobic species in our study reduced their overall activity during periods with new moon. If these results extend to artificial light, a loss of dark nights could curtail the amount of time some species invest into foraging and other important activities. Strong responses to artificial light have already been observed in nocturnal mammals (Prugh and Golden 2014). For example, the common spiny mouse (*Acomys cahirinus*) shows a clear reduction in the overall activity and time for foraging when exposed to artificial light (Rotics et al. 2011). The constant reduction of activity, for example due to permanent human light sources, may, affect individuals, populations and even communities. However, predicting fitness consequences of artificial light based on responses to lunar phases is challenging. Seemingly indifferent species without adaptations to changing nocturnal light conditions may not be impacted at all or could bear the brunt of brighter nights resulting from canopy loss and light pollution if they are made vulnerable by increased visibility. Species that change their overall activity level in response to nocturnal illumination may be more strongly impacted than species that can maintain their level of activity by adjusting its timing. Along those lines, lunar phobic species could be expected to cope better with artificial light if they follow a cathemeral diel activity pattern as this is indicative of behavioral plasticity that may be advantageous in a changing world ((temperature changes, artificial light; Cox and Gaston 2023). However, in tropical regions cathemerality is less reportedly less common than in higher latitudes (Bennie et al. 2014).

Our research describes responses to moonlight on the forest floor. It would be interesting for future studies to examine responses in the canopy of tropical forest, where lunar illumination likely has more pronounced effects on animal behavior. It is also worthwhile to extend research into the effects of moonlight and artificial light to birds, another prominent class of both ground and canopy-dwelling species in tropical forests.

## Acknowledgements

This work was made possible by the Tropical Ecology Assessment and Monitoring (TEAM) Network, a collaboration between Conservation International, the Smithsonian Tropical Research Institute, and the Wildlife Conservation Society that is partially funded by these organizations, the Gordon and Betty Moore Foundation, and other donors. We acknowledge the effort of all TEAM site managers and collaborators who helped collect data as well as Wildlife Insight for data processing and availability. This study received partial funding from the Research Council of Norway (Grant nr. NFR 301075).

## Data availability statement

The data and code for performing the analyses described in this article are available at: https://github.com/richbi/TropicalMoon.

## Supplementary information

Appendix S1

