## Supplementary Information for "The moon’s influence on the activity of tropical forest mammals"

### Appendix S1

**Article:** The moon's influence on the activity of tropical forest mammals

**Journal:** Ecology

**Authors:** Richard Bischof, Andrea F. Vallejo-Vargas, Asunción Semper-Pascual, Simon D. Schowanek, Lydia Beaudrot, Daniel Turek, Patrick A. Jansen[4], Francesco Rovero, Steig E. Johnson, Marcela Guimarães Moreira Lima, Fernanda Santos, Eustrate Uzabaho, Santiago Espinosa, Jorge A. Ahumada, Robert Bitariho, Julia Salvador, Badru Mugerwa, Moses N. Sainge, Douglas Sheil

**List of contents:**

Tables S1, S2, and S3

Figures S1, S2, and S3

**Table S1.** Designation of forest mammals in the Afrotropics to diel and lunar categories based on multinomial logistic regression and ternary classification. Also shown are the number of observations (15-minute intervals with at least one photographic detection of a species at distinct camera trap locations).

| | | **Designation** | | **Number of observations** | | | | | | |
|---|---|---|---|---|---|---|---|---|---|---|
| **Species** | **Order** | **Diel category** | **Lunar category** | **Full moon** | **Tran-sition** | **New moon** | **Night** | **Twiligh t** | **Day** | **Total** |
| *Atherurus africanus* | Rodentia | nocturnal | lunarphobic | 44 | 244 | 664 | 952 | 109 | 55 | 1116 |
| *Atilax paludinosus* | Carnivora | cathemeral | indifferent | 9 | 19 | 29 | 57 | 33 | 99 | 189 |
| *Bdeogale crassicauda* | Carnivora | noct.-crep. | indifferent | 467 | 899 | 1339 | 2705 | 499 | 9 | 3213 |
| *Bdeogale nigripes* | Carnivora | nocturnal | indifferent | 22 | 42 | 88 | 152 | 10 | 13 | 175 |
| *Canis adustus* | Carnivora | noct.-crep. | indifferent | 17 | 18 | 20 | 55 | 34 | 41 | 130 |
| *Caracal aurata* | Carnivora | cathemeral | indifferent | 20 | 38 | 72 | 130 | 51 | 178 | 359 |
| *Cephalophus callipygus* | Artiodactyla | diurn.-crep. | indifferent | 151 | 241 | 384 | 776 | 858 | 9197 | 10831 |
| *Cephalophus dorsalis* | Artiodactyla | noct.-crep. | indifferent | 329 | 547 | 831 | 1707 | 297 | 237 | 2241 |
| *Cephalophus harveyi* | Artiodactyla | diurn.-crep. | indifferent | 82 | 133 | 183 | 398 | 334 | 3197 | 3929 |
| *Cephalophus nigrifrons* | Artiodactyla | diurn.-crep. | indifferent | 180 | 330 | 527 | 1037 | 546 | 5708 | 7291 |
| *Cephalophus silvicultor* | Artiodactyla | noct.-crep. | indifferent | 516 | 925 | 1134 | 2575 | 962 | 855 | 4392 |
| *Cephalophus spadix* | Artiodactyla | cathemeral | indifferent | 28 | 48 | 61 | 137 | 71 | 300 | 508 |
| *Civettictis civetta* | Carnivora | noct.-crep. | indifferent | 9 | 16 | 14 | 39 | 19 | 3 | 61 |
| *Cricetomys emini* | Rodentia | nocturnal | lunarphobic | 120 | 489 | 1663 | 2272 | 229 | 74 | 2575 |
| *Cricetomys gambianus* | Rodentia | noct.-crep. | indifferent | 387 | 1018 | 2784 | 4189 | 626 | 43 | 4858 |
| *Cryptoprocta ferox* | Carnivora | cathemeral | indifferent | 17 | 18 | 37 | 72 | 40 | 66 | 178 |
| *Dendrohyrax arboreus* | Hyracoidea | noct.-crep. | indifferent | 10 | 16 | 33 | 59 | 11 | 3 | 73 |
| *Dendrohyrax validus* | Hyracoidea | noct.-crep. | indifferent | 46 | 101 | 145 | 292 | 60 | 11 | 363 |
| *Eliurus tanala* | Rodentia | noct.-crep. | lunarphobic | 6 | 26 | 64 | 96 | 18 | 24 | 138 |
| *Fossa fossana* | Carnivora | noct.-crep. | indifferent | 78 | 171 | 228 | 477 | 132 | 19 | 628 |
| *Galidictis fasciata* | Carnivora | noct.-crep. | indifferent | 7 | 6 | 12 | 25 | 5 | 0 | 30 |
| *Genetta servalina* | Carnivora | noct.-crep. | indifferent | 108 | 212 | 395 | 715 | 173 | 45 | 933 |
| *Gorilla gorilla* | Primates | diurnal | indifferent | 3 | 11 | 18 | 32 | 11 | 387 | 430 |
| *Hyemoschus aquaticus* | Artiodactyla | nocturnal | indifferent | 22 | 33 | 26 | 81 | 3 | 2 | 86 |
| *Loxodonta africana* | Proboscidea | cathemeral | indifferent | 83 | 154 | 181 | 418 | 129 | 550 | 1097 |

| | | | | | | | | | | |
|---|---|---|---|---|---|---|---|---|---|---|
| *Nandinia binotata* | Carnivora | noct.-crep. | indifferent | 23 | 44 | 67 | 134 | 25 | 3 | 162 |
| *Nesomys rufus* | Rodentia | diurn.-crep. | lunar phobic | 0 | 4 | 30 | 34 | 84 | 334 | 452 |
| *Nesotragus moschatus* | Artiodactyla | cathemeral | indifferent | 155 | 180 | 174 | 509 | 353 | 455 | 1317 |
| *Orycteropus afer* | Tubulidentata | nocturnal | indifferent | 5 | 11 | 17 | 33 | 1 | 1 | 35 |
| *Pan troglodytes* | Primates | diurnal | NA | 5 | 16 | 8 | 29 | 73 | 965 | 1067 |
| *Panthera pardus* | Carnivora | cathemeral | indifferent | 12 | 20 | 19 | 51 | 28 | 73 | 152 |
| *Petrodromus tetradactylus* | Macroscelidea | noct.-crep. | lunar philic | 37 | 38 | 23 | 98 | 40 | 2 | 140 |
| *Phataginus tricuspis* | Pholidota | noct.-crep. | indifferent | 9 | 22 | 35 | 66 | 11 | 2 | 79 |
| *Philantomba monticola* | Artiodactyla | diurnal | indifferent | 178 | 358 | 518 | 1054 | 1014 | 14065 | 16133 |
| *Potamochoerus larvatus* | Artiodactyla | cathemeral | indifferent | 40 | 65 | 98 | 203 | 84 | 193 | 480 |
| *Potamochoerus porcus* | Artiodactyla | noct.-crep. | indifferent | 100 | 117 | 178 | 395 | 131 | 218 | 744 |
| *Protoxerus stangeri* | Rodentia | diurnal | indifferent | 10 | 36 | 49 | 95 | 20 | 483 | 598 |
| *Syncerus caffer* | Artiodactyla | cathemeral | indifferent | 65 | 73 | 105 | 243 | 91 | 328 | 662 |
| *Tenrec ecaudatus* | Afrosoricida | noct.-crep. | indifferent | 7 | 10 | 22 | 39 | 10 | 9 | 58 |
| *Tragelaphus scriptus* | Artiodactyla | cathemeral | indifferent | 246 | 418 | 522 | 1186 | 431 | 1633 | 3250 |

**Table S2.** Designation of forest mammals in the Neotropics to diel and lunar categories based on multinomial logistic regression and ternary classification. Also shown are the number of observations (15-minute intervals with at least one photographic detection at distinct camera trap locations).

| | | **Designation** | | **Number of observations** | | | | | | |
|---|---|---|---|---|---|---|---|---|---|---|
| **Species** | **Order** | **Diel category** | **Lunar category** | **Full moon** | **Tran-sition** | **New moon** | **Night** | **Twilight** | **Day** | **Total** |
| *Cuniculus paca* | Rodentia | nocturnal | lunar phobic | 376 | 1565 | 4434 | 6375 | 684 | 56 | 7115 |
| *Dasyprocta fuliginosa* | Rodentia | diurnal | indifferent | 3 | 7 | 19 | 29 | 142 | 2806 | 2977 |
| *Dasyprocta leporina* | Rodentia | diurnal | indifferent | 7 | 19 | 21 | 47 | 388 | 3940 | 4375 |
| *Dasyprocta punctata* | Rodentia | diurn.-crep. | indifferent | 37 | 69 | 81 | 187 | 1381 | 11750 | 13318 |
| *Dasypus kappleri* | Cingulata | nocturnal | lunar phobic | 55 | 162 | 579 | 796 | 50 | 2 | 848 |
| *Dasypus novemcinctus* | Cingulata | nocturnal | lunar phobic | 160 | 522 | 1909 | 2591 | 226 | 60 | 2877 |
| *Didelphis marsupialis* | Didelphimorphia | noct.-crep. | indifferent | 283 | 421 | 707 | 1411 | 224 | 35 | 1670 |
| *Hoplomys gymnurus* | Rodentia | nocturnal | lunar phobic | 1 | 5 | 34 | 40 | 5 | 0 | 45 |
| *Hylaeamys megacephalus* | Rodentia | noct.-crep. | lunar phobic | 1 | 7 | 25 | 33 | 5 | 0 | 38 |
| *Leopardus pardalis* | Carnivora | noct.-crep. | indifferent | 138 | 241 | 360 | 739 | 210 | 298 | 1247 |
| *Leopardus wiedii* | Carnivora | noct.-crep. | indifferent | 27 | 33 | 52 | 112 | 18 | 18 | 148 |
| *Mazama americana* | Artiodactyla | noct.-crep. | indifferent | 532 | 770 | 940 | 2242 | 795 | 1255 | 4292 |
| *Mazama nemorivaga* | Artiodactyla | diurnal | indifferent | 18 | 32 | 41 | 91 | 41 | 1094 | 1226 |
| *Mazama temama* | Artiodactyla | cathemeral | indifferent | 61 | 135 | 153 | 349 | 104 | 603 | 1056 |
| *Metachirus nudicaudatus* | Didelphimorphia | nocturnal | indifferent | 56 | 161 | 487 | 704 | 75 | 16 | 795 |
| *Myoprocta acouchy* | Rodentia | diurn.-crep. | indifferent | 4 | 10 | 19 | 33 | 495 | 1751 | 2279 |
| *Myoprocta pratti* | Rodentia | diurn.-crep. | indifferent | 7 | 11 | 24 | 42 | 495 | 2302 | 2839 |
| *Neacomys spinosus* | Rodentia | nocturnal | indifferent | 3 | 8 | 18 | 29 | 3 | 0 | 32 |
| *Odocoileus virginianus* | Artiodactyla | diurn.-crep. | indifferent | 16 | 19 | 32 | 67 | 37 | 252 | 356 |
| *Panthera onca* | Carnivora | cathemeral | indifferent | 17 | 30 | 35 | 82 | 17 | 133 | 232 |
| *Pecari tajacu* | Artiodactyla | diurn.-crep. | indifferent | 214 | 309 | 450 | 973 | 634 | 5398 | 7005 |
| *Philander opossum* | Didelphimorphia | nocturnal | lunar phobic | 26 | 80 | 224 | 330 | 27 | 2 | 359 |
| *Priodontes maximus* | Cingulata | nocturnal | indifferent | 13 | 28 | 103 | 144 | 15 | 10 | 169 |
| *Procyon cancrivorus* | Carnivora | noct.-crep. | indifferent | 8 | 26 | 33 | 67 | 15 | 2 | 84 |
| *Proechimys brevicauda* | Rodentia | noct.-crep. | lunar phobic | 84 | 292 | 887 | 1263 | 186 | 44 | 1493 |

| | | | | | | | | | | |
|---|---|---|---|---|---|---|---|---|---|---|
| *Proechimys guyannensis* | Rodentia | nocturnal | indifferent | 12 | 35 | 85 | 132 | 15 | 0 | 147 |
| *Proechimys semispinosus* | Rodentia | noct.-crep. | indifferent | 57 | 153 | 316 | 526 | 83 | 0 | 609 |
| *Puma concolor* | Carnivora | cathemeral | indifferent | 21 | 39 | 44 | 104 | 39 | 184 | 327 |
| *Sylvilagus brasiliensis* | Lagomorpha | crepuscular | lunar philic | 26 | 11 | 6 | 43 | 54 | 4 | 101 |
| *Tamandua mexicana* | Pilosa | cathemeral | indifferent | 17 | 16 | 43 | 76 | 19 | 63 | 158 |
| *Tamandua tetradactyla* | Pilosa | noct.-crep. | indifferent | 7 | 8 | 32 | 47 | 12 | 17 | 76 |
| *Tapirus bairdii* | Perissodactyla | noct.-crep. | indifferent | 39 | 83 | 103 | 225 | 60 | 89 | 374 |
| *Tapirus terrestris* | Perissodactyla | noct.-crep. | indifferent | 211 | 297 | 423 | 931 | 271 | 277 | 1479 |
| *Tayassu pecari* | Artiodactyla | diurnal | lunar philic | 22 | 17 | 10 | 49 | 28 | 612 | 689 |
| *Tylomys watsoni* | Rodentia | nocturnal | lunar phobic | 3 | 5 | 25 | 33 | 1 | 0 | 34 |

**Table S3.** Designation of forest mammals in the Indomalayan tropics to diel and lunar categories based on multinomial logistic regression and ternary classification. Also shown are the number of observations (15-minute intervals with at least one photographic detection at distinct camera trap locations).

| | | Designation | | Number of observations | | | | | | |
|---|---|---|---|---|---|---|---|---|---|---|
| **Species** | **Order** | **Diel category** | **Lunar category** | **Full moon** | **Tran-sition** | **New moon** | **Night** | **Twiligh t** | **Day** | **Total** |
| *Atherurus macrourus* | Rodentia | nocturnal | indifferent | 29 | 75 | 211 | 315 | 32 | 6 | 353 |
| *Echinosorex gymnura* | Eulipotyphla | nocturnal | indifferent | 8 | 10 | 22 | 40 | 5 | 0 | 45 |
| *Hemigalus derbyanus* | Carnivora | nocturnal | indifferent | 7 | 18 | 30 | 55 | 5 | 1 | 61 |
| *Hystrix brachyura* | Rodentia | nocturnal | lunar phobic | 28 | 100 | 315 | 443 | 39 | 7 | 489 |
| *Leopoldamys sabanus* | Rodentia | nocturnal | lunar phobic | 19 | 21 | 127 | 167 | 18 | 3 | 188 |
| *Macaca nemestrina* | Primates | diurnal | indifferent | 11 | 14 | 31 | 56 | 30 | 3192 | 3278 |
| *Muntiacus muntjak* | Artiodactyla | diurn.-crep. | indifferent | 53 | 74 | 83 | 210 | 157 | 949 | 1316 |
| *Paradoxurus hermaphroditus* | Carnivora | noct.-crep. | indifferent | 10 | 16 | 24 | 50 | 10 | 3 | 63 |
| *Prionailurus bengalensis* | Carnivora | noct.-crep. | indifferent | 7 | 14 | 15 | 36 | 9 | 8 | 53 |
| *Rusa unicolor* | Artiodactyla | cathemeral | indifferent | 6 | 14 | 13 | 33 | 15 | 88 | 136 |
| *Sus scrofa* | Artiodactyla | cathemeral | indifferent | 83 | 169 | 286 | 538 | 278 | 1402 | 2218 |
| *Tapirus indicus* | Perissodactyla | noct.-crep. | indifferent | 28 | 44 | 61 | 133 | 64 | 54 | 251 |
| *Tragulus sp.* | Artiodactyla | diurn.-crep. | indifferent | 37 | 47 | 52 | 136 | 202 | 554 | 892 |

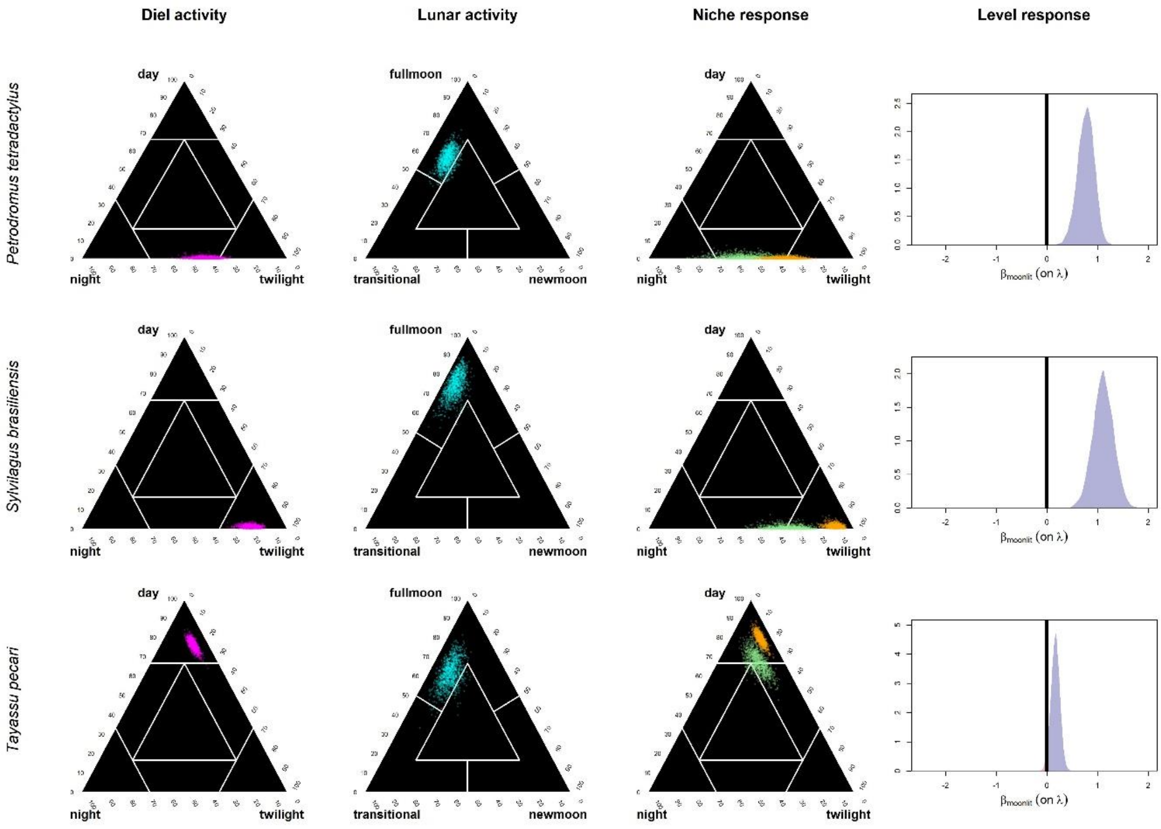


**Figure S1**. Overview of ternary classifications for diel and lunar activity (columns 1-2) and responses to lunar phases (column 3 and 4) for three mammal species identified as lunar philic. Column 1: ternary plots of diel activity posteriors. Column 2: ternary plot of lunar activity posteriors. Column 3: ternary plots showing difference of diel activity (potential temporal niche shifting) between periods with full moon (green) and without moonlit nights (orange). Column 4: posterior distribution of the difference between overall activity (related to the number of photographic detection events) during periods with vs. without full moon. Negative (red) and positive (blue) values indicate a reduction or an increase, respectively, in overall activity during periods (multiple 24-hour periods, see Fig. 1F in the main text) with full moon.

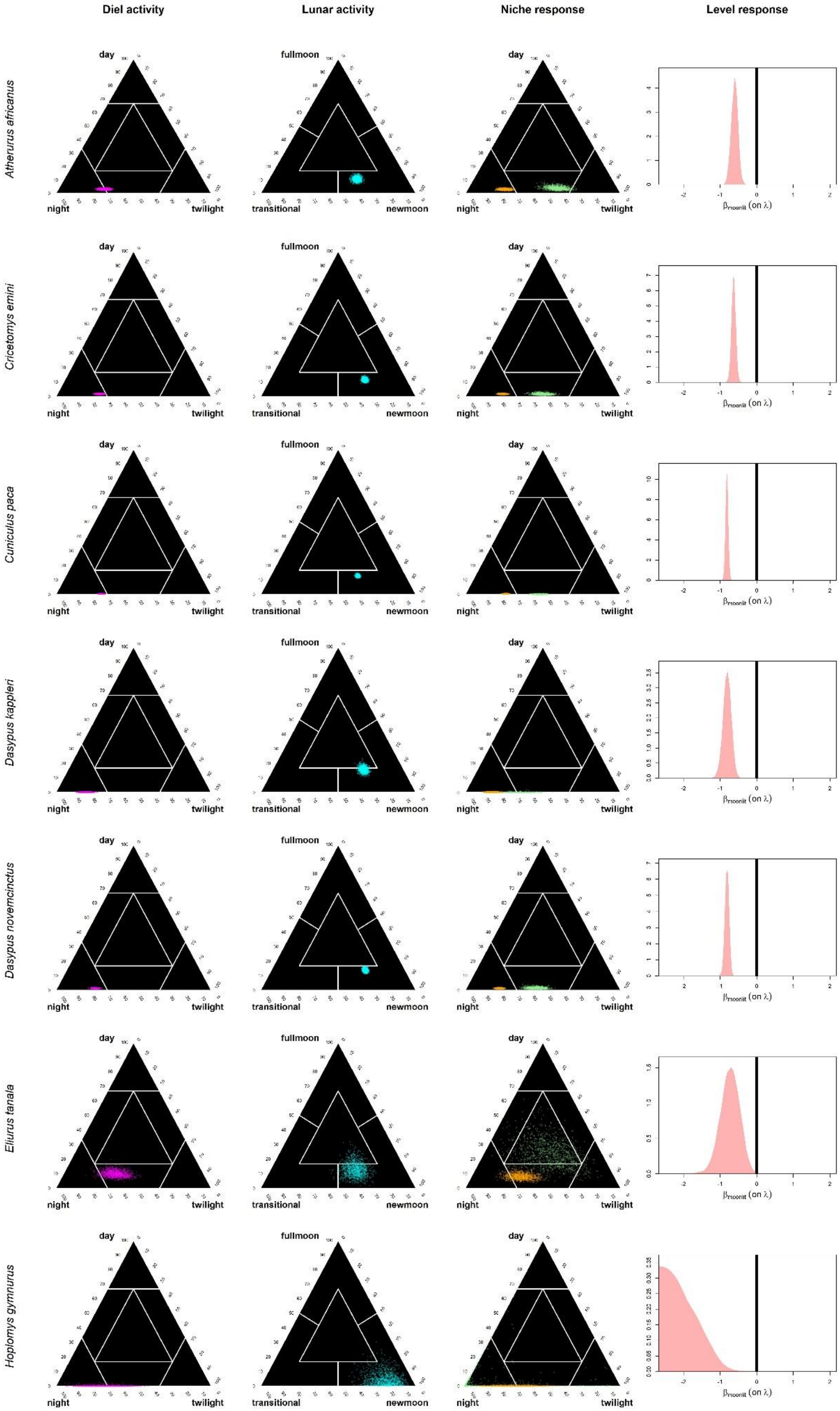
Diel activity
Lunar activity
Niche response
Level response
Atherurus africanus
Cricetomys emini
Cuniculus paca
Dasypus kappleri
Dasypus novemcinctus
Eliurus tanala
Hoplomys gymnurus
day
night
twilight
fullmoon
transitional
newmoon
βmoonlit (on λ)

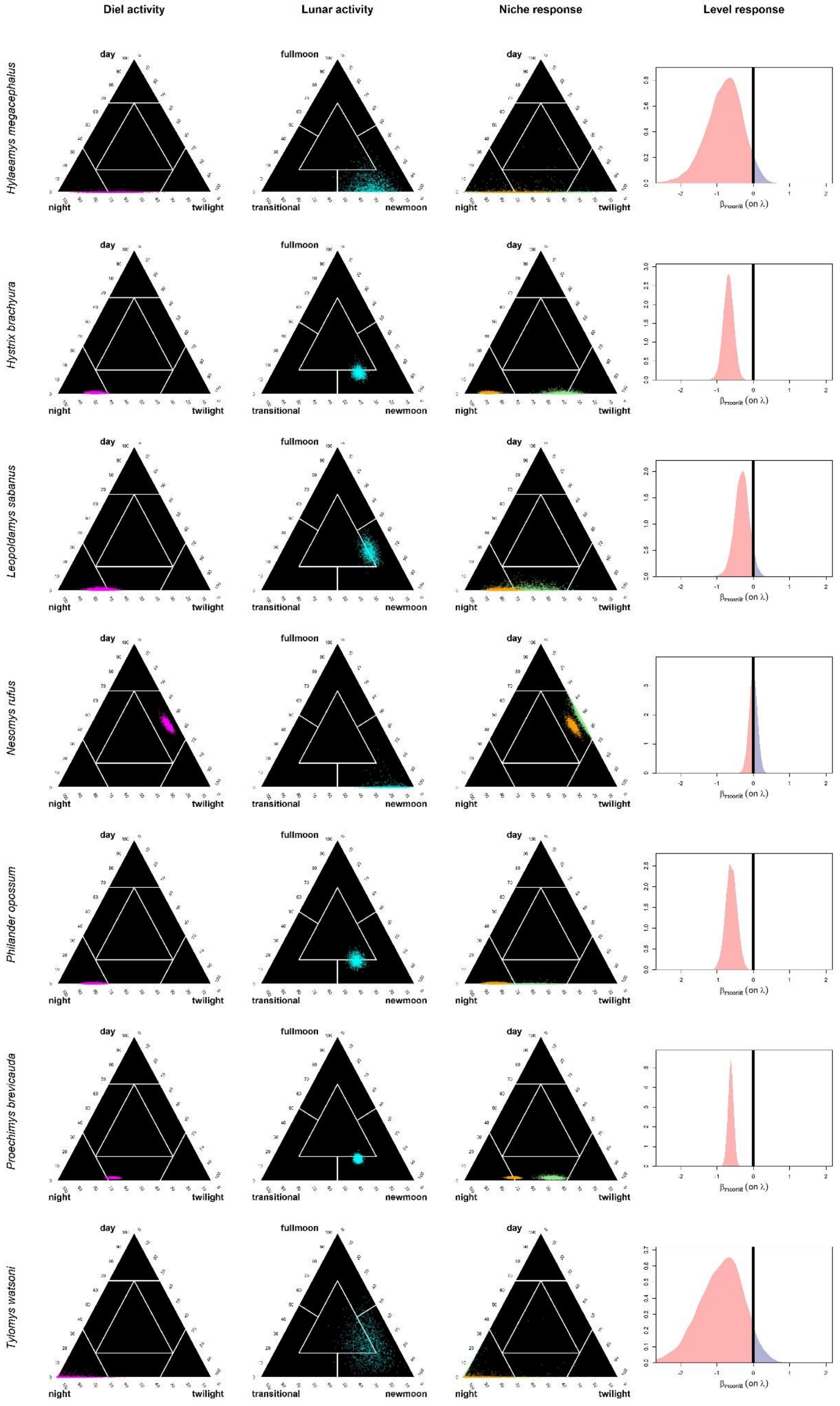
Diel activity
Lunar activity
Niche response
Level response
Hylaeamys megacephalus
Hystrix brachyura
Leopoldamys sabanus
Nesomys rufus
Philander opossum
Proechimys brevicauda
Tylomys watsoni
day
night
twilight
fullmoon
transitional
newmoon
β moonlit (on λ)

**Figure S2**. Overview of ternary classifications for diel and lunar activity (columns 1-2) and responses to lunar phases (column 3 and 4) for 14 mammal species identified as lunar phobic. Column 1: ternary plots of diel activity posteriors. Column 2: ternary plot of lunar activity posteriors. Column 3: ternary plots showing difference of diel activity (potential temporal niche shifting) between periods with full moon (green) and without moonlit nights (orange). Column 4: posterior distribution of the difference between overall activity (related to the number of photographic detection events) during periods with vs. without full moon. Negative (red) and positive (blue) values indicate a reduction or an increase, respectively, in overall activity during periods (multiple 24-hour periods, see Fig. 1F in the main text) with full moon.
